# Dyrk1a is required for craniofacial development in *Xenopus laevis*

**DOI:** 10.1101/2024.01.13.575394

**Authors:** H. Katherine Johnson, Stacy E Wahl, Fatmata Sesay, Larisa Litovchick, Amanda JG Dickinson

**Affiliations:** Department of Biology, Virginia Commonwealth University, Richmond, Virginia, USA; Department of Internal Medicine, Division of Hematology, Oncology and Palliative Care, Virginia Commonwealth University, Richmond, Virginia, USA; Massey Comprehensive Cancer Center, Richmond, Virginia, USA

**Author notes:** School of Medicine Office of Faculty Affairs and Faculty Development, Virginia Commonwealth University, Richmond, Virginia, USA. **Corresponding author:** Amanda Dickinson.

**Keywords:** DYRK1A, craniofacial, *Xenopus laevis*

## Abstract

Loss of function mutations in the dual specificity tyrosine-phosphorylation-regulated kinase 1A (DYRK1A) gene are associated with craniofacial malformations in humans. Here we characterized the effects of deficient DYRK1A in craniofacial development using a developmental model, *Xenopus laevis*. *Dyrk1a* mRNA and protein was expressed throughout the developing head and was enriched in the branchial arches which contribute to the face and jaw. Consistently, reduced Dyrk1a function, using *dyrk1a* morpholinos and pharmacological inhibitors, resulted in orofacial malformations including hypotelorism, altered mouth shape, slanted eyes, and narrower face accompanied by smaller jaw cartilage and muscle. Inhibition of Dyrk1a function resulted in misexpression of key craniofacial regulators including transcription factors and members of the retinoic acid signaling pathway. Two such regulators, *sox9* and *pax3* are required for neural crest development and their decreased expression corresponds with smaller neural crest domains within the branchial arches. Finally, we determined that the smaller size of the faces, jaw elements and neural crest domains in embryos deficient in Dyrk1a could be explained by increased cell death and decreased proliferation. This study is the first to provide insight into why craniofacial birth defects might arise in humans with DYRK1A mutations.

## Introduction

DYRK1A (<u>D</u>ual-<u>s</u>pecificity Tyrosine (<u>Y</u>) <u>R</u>egulated <u>K</u>inase) is a multifunctional kinase that phosphorylates protein substrates involved in a wide range of cellular process such as cell cycle, transcription, signaling, and apoptosis (reviewed in (Ananthapadmanabhan et al., 2023; Atas-Ozcan et al., 2021; Fernandez-Martinez et al., 2015)). Therefore, it is not surprising that proper dosage of this protein is critical for a variety of developmental processes. Indeed, in humans, haploinsufficiency of *DYRK1A* is associated with DYRK1A syndrome (aka DYRK1A haploinsufficiency syndrome, Intellectual developmental disorder, autosomal dominant 7, MRD7, OMIM # 614104). This developmental disorder is characterized by intellectual disability, microcephaly, short stature and malformations affecting the kidney and craniofacial structures (Blackburn et al., 2019; Ji et al., 2015; van Bon BWM, 2015; Widowati et al., 2018). Consistently, work in vertebrate animal models has demonstrated that decreasing Dyrk1a affects the developing brain and pronephros in line with phenotypes in humans (Blackburn et al., 2019; Fotaki et al., 2002; Kim et al., 2017; Willsey et al., 2020). While humans with DYRK1A syndrome have a myriad of craniofacial malformations such as retrognathia, and differences in the nose, upper lip and philtrum, there has not been any investigations that specifically address why DYRK1A haploinsufficiency results in these craniofacial differences. Therefore, the broad goals of this study is to 1) investigate the consequences of decreased DYRK1A on craniofacial development and 2) begin to dissect the mechanisms by which craniofacial malformations occur when Dyrk1a dosage is reduced. We do this using an effective developmental model, *Xenopus laevis,* where Dyrk1a function can be titrated and embryos can easily be evaluated throughout all stages of early development.

DYRK1A has a multitude of functions and diverse substrates, therefore it is a challenging but important gene to understand during development. Notably*, DYRK1A* is also within the Down Syndrome critical region. Increased dosage of this gene is thought to be one of the contributors to a range of developmental malformations associated with this syndrome including craniofacial differences (Atas-Ozcan et al., 2021; Stringer et al., 2017). Although there is limited research specifically investigating the impact of DYRK1A gene loss on craniofacial development in whole animal models, we do know that the protein is important in the context of Down Syndrome and DYRK1A overexpression models. For example, in mouse models of Down Syndrome, the use of DYRK1A inhibitors or genetic knockout of DYRK1A has been shown to mitigate skull and jaw malformations in such models (McElyea et al., 2016; Redhead et al., 2023). Thus, decreased, or increased dosage of DYRK1A is detrimental to craniofacial development indicating that Dyrk1a plays a crucial role in the development of the face.

In all vertebrates, the earliest stages of craniofacial development are very similar. For example, the presumptive facial region is influenced by cranial neural crest cells (CNCs), which originate next to the neural tube and migrate into the head (Cordero et al., 2011; Minoux and Rijli, 2010). These CNCs as well as facial ectoderm, endoderm, and mesoderm contribute to the facial prominences that form around the embryonic mouth. The facial prominences then grow, fuse and differentiate into specific tissues that form the midface, palate and jaws (Bush and Jiang, 2012; Dudas et al., 2007; Suzuki et al., 2016). The development of the CNCs, growth of the facial prominences, and differentiation of jaw elements are orchestrated by a similar set of genes and signaling pathways across vertebrates (Dickinson, 2016; Francis-West et al., 2003; Helms et al., 2005; Jacox et al., 2014; Kennedy and Dickinson, 2012; Van Otterloo et al., 2016; Wahl et al., 2015).

We have utilized the vertebrate developmental model, *Xenopus laevis,* to fill a major gap and study the effects of decreased Dyrk1a on craniofacial development. This model has been established as an effective tool for studying how the face forms (Dickinson, 2016; Dickinson, 2023; Dubey and Saint-Jeannet, 2017; Kennedy and Dickinson, 2012, 2014; Sullivan and Levin, 2016; Vandenberg et al., 2012; Wyatt et al., 2020). Moreover, deficiency or mutations in the same genes that cause craniofacial abnormalities in humans also cause similar defects in *Xenopus* (Adams et al., 2016; Bajpai et al., 2010; Devotta et al., 2016; Dubey et al., 2018; Dubey and Saint-Jeannet, 2017; Greenberg et al., 2019; Hwang et al., 2019; Lasser et al., 2019; Mills et al., 2019; Mohammadparast and Chang, 2022; Parast et al., 2023; Tahir et al., 2014; Wahl et al., 2015; Wyatt et al., 2020). In the present study, we demonstrate that decreasing Dyrk1a function in *X. laevis* results in craniofacial malformations that correlate with changes in the expression of craniofacial regulators as well as increased apoptosis in the developing head.

## Methods

### Obtaining Xenopus laevis embryos

*Xenopus laevis* embryos were obtained using standard procedures (Sive et al., 2000) approved by the VCU Institutional Animal Care and Use Committee (IACUC protocol number AD20261). Embryos were staged according to Nieuwkoop and Faber (Nieuwkoop and Faber, 1967; Nieuwkoop and Faber, 1994). Stages are also reported as hours post fertilization at 23°C for better comparisons across vertebrates.

### Morpholino Knockdown

To knockdown gene function we used antisense oligonucleotides stabilized with morpholino rings (Morpholinos (MOs) GeneTools). *X. laevis* is tetraploid and thus has two homeologs of each gene (denoted S and L). We used a previously validated translation blocking Dyrk1 MO that targets both homeologs of this gene (Xenbase.org). A standard control MO that does not target any Xenopus sequence was utilized as a negative control. MOs were labeled with fluorescein which allowed separation of un-injected individuals by 24 hours of development. Embryos were injected with a Femtojet injection system using standard practices (Sive et al., 2010). Successfully injected embryos, as indicated by green fluorescence and being alive, were sorted under a stereoscope and raised at 15°C in frog embryo media (0.1X Modified Barth’s Saline (MBS)). MOs were microinjected using an Eppendorf Femtojet microinjector and a Zeiss Discovery V8 stereoscope. Embryos were placed in a dish lined with nylon Spectra mesh (1,000 um opening and 1350 um thickness) at the bottom to hold embryos in place and filled with 3% Ficoll 400 (Fisher, BP525) dissolved in frog embryo media. After injections, embryos were placed into 10 cm culture dishes filled with frog embryo media and maintained in a 15°C incubator. Media was changed daily, and dead embryos were removed. Three biological replicates were performed where embryos were generated from different parents on different days.

### DYRK1A pharmacological antagonist treatments

Embryos were immersed in 25uM INDY (Tocris, 4997) or 50uM Harmine (Sigma-Aldrich, SMB00461) with 0.1% DMSO in 4 ml of frog embryo media (0.1X MBS) in 6 well culture dishes. Controls were immersed in 0.1% DMSO alone in the same volume and in the same dish. Equal numbers of embryos (15 embryos), were placed in each well and 2 technical replicates (embryos from the same parents on the same day) of each performed. At least 2 biological replicates (embryos from different parents on a different day) were performed. Embryos were treated at stage 22-24 (∼24 hpf) until stage 32-33 at 15°C and then processed for activity assays, qRT-PCR, staining or washed out and cultured until the desired stage in frog embryo media in 10 cm petri dishes.

### DYRK1A in vitro kinase assay

Embryos exposed to Dyrk1a inhibitors or injected with dyrk1a MOs were placed in microcentrifuge tube at stage 26 and washed in 50mM Tris buffer. The liquid was removed and tubes stored at -80C. Purified recombinant GST-LIN52 was incubated with embryo extracts in the presence of ATP then phosphorylated LIN52-S28, total GST-LIN52, DYRK1A and loading control proteins were assessed by western blotting as previously described (Iness et al., 2019; Menon et al., 2019).

### Imaging the faces of Xenopus laevis

At stage 43, embryos were anesthetized in 1% tricaine for 10 minutes and then fixed in 4% paraformaldehyde overnight at 4°C. The heads were removed to effectively view and image the face and head. A No. 15 scalpel (VWR, 82029-856) and Dumont No. 5 forceps (Fisher, NC9404145) were used to make two cuts to isolate the head: first-at the posterior end of the gut and then second caudal to the gills. Isolated heads were mounted in small holes or depressions in either agarose or clay-lined dishes containing Phosphate Buffered Saline with 0.1% Tween (PBT). The faces were imaged using a Zeiss stereoscope fitted with an Axiovision digital camera (Zeiss).

### Facial Measurements and Statistics

Measurements of intercanthal distance, the distance from the inside of each eye across the midface was measured using Zeiss Zen Blue software. Data was imported into Excel to create stacked bar graphs. 60 embryos in 2-3 biological replicates were quantified for each treatment group. SigmaPlot was utilized to perform statistical analyses and to create box and whisker plots. A multiple-group analysis was performed where it was first analyzed for normality and equal variance. An ANOVA or ANOVA on Ranks (Kruskal Wallis test) was performed to determine if there were statistical differences across all treatment groups. This was then followed by a Tukey posthoc test to compare two groups. Significance was deemed if p-values were lower than 0.05.

### Immunofluorescence and Quantification of CC3 and PH3

To perform all fluorescent labeling, embryos were fixed in 4% paraformaldehyde. To create transverse sections through the oral cavity, fixed embryos were immersed in 5% low melt agarose, and 200 micron sections created with a vibratome (Leica). The sections or whole embryos were incubated in blocking buffer (1% Triton-X, 1% normal calf serum, 1% BSA in PBT) overnight at 4°C. This was followed by primary antibody incubation (anti-DYRK1A, 1:100, Bethyl Laboratories, A303-802A), Cleaved Caspase-3 (CC3, 1:1000, Cell Signaling Technology, 9661S), phospho-histone H3 (PH3, 1:100, Sigma, 06-570), collagen II (Developmental Studies Hybridoma Bank, 1:25), and 12/101 (Developmental Studies Hybridoma Bank, 1:25) for 2-4 days and then washed 3 times in 1 hour at room temperature. Sections or whole embryos were then incubated in secondary antibody (1:500, goat-anti mouse Alexa Fluor 488 or 568) combined with 2 drops of NucBlue in 1 ml PBT (Invitrogen, RF37606), for 1-2 days. For double labeling, sections or embryos were labeled with the first primary and corresponding secondary antibody followed by the second primary and corresponding secondary antibodies. Since individual cells could not be discerned in the CC3 labeling, the histogram function in Photoshop was used to assess fluorescence. The mean intensity of the fluorescence was determined for the face, excluding the brain and eyes, and normalized to area. Individual PH3 positive cells were counted in the face, excluding the brain and eyes, and also normalized to area. SigmaPlot was used to perform statistical analysis consisting of TTESTs for 2 groups or ANOVAs followed by Tukey posthoc test for 3 groups.

### Cartilage Labeling

Cartilages were labeled with Alcian Blue as described previously. Tadpoles were fixed in Bouin’s fixative overnight at 4°C and then washed in 70% ethanol until clear of any yellow color. They were then immersed in Alcian Blue mixture (0.1mg/ml Alcian Blue in 1 part acetic acid: 4 parts ethanol) for 3 days at room temperature. Specimens were washed in 1% HCL in 70% ethanol for 1–2 days followed by another fixation in 4% paraformaldehyde (2 hrs). To remove pigment, tadpoles were immersed in 1.5% hydrogen peroxide and 5% formamide in saline sodium citrate solution and placed on a lightbox until pigment could no longer be seen. After three washes in PBT, the embryos were incubated in 2% potassium hydroxide (2 times for 30 minutes). Finally, the embryos were cleared in a series of 2% potassium hydroxide to 100% glycerol solutions (50:50, 40:60, 20:80, 30 minutes each). Tadpoles were then imaged in 100% glycerol.

### In situ hybridization

DIG-labeled RNA probes for *dyrk1a* were prepared from a plasmid containing a partial, sequence of Xenopus laevis dyrk1a.S (IMAGE Clone ID: 7010948, now available at Horizon, MXL1736-202713005). Due to the high degree of similarity, a probe made from this sequence was predicted to bind to both *dyrk1a.L* and *dyrk1a.S*. The plasmid was linearized with Sal1 (antisense) or Not1 (sense), gel purified and transcribed with DIG RNA labeling mix with either T7 RNA polymerase (NEB, M0251S; antisense probe), or SP6 RNA polymerase (NEB, M0207S; sense probe). Embryos used for in situ hybridization were fixed in 4% PFA, dehydrated in a methanol series, and stored for 1–10 days at −20C. In situ hybridization was performed as described (Sive et al., 2000), omitting the proteinase K treatment.

### *sox10*-GFP transgenic assessment

*sox10*-GFP transgenic adult *Xenopus laevis* (*Xla.Tg(sox10:GFP)Jpsj,* RRID: NXR_0108) originating in the Saint-Jeannet lab were obtained from Marine Biological Laboratory National Xenopus Resource (Alkobtawi et al., 2018; Bowes et al., 2008; Ossipova et al., 2018). Embryos were obtained by natural mating and then exposed to INDY as described above. Embryos were fixed in 4% paraformaldehyde for 24 hours and washed in PBT and then assessed on a Nikon C2 confocal microscope.

### qRT-PCR of craniofacial regulators

Craniofacial gene expression was assessed using quantitative reverse transcription PCR (RT-qPCR). RNA was extracted from the heads of embryos at stage 32 using TRIzol extraction following the provided instructions followed by lithium chloride precipitation. One-step quantitative RT-PCR was performed using the extracted RNA, and both reverse transcription and PCR were performed using the Luna One-Step Universal RT-qPCR kit (NEB, E3005S) and carried out on the Mic qPCR Machine (BioMolecular Systems) using the primer sequences included in Supplemental Table 1. Using the delta-delta CT method, the relative fold change between control and experimental groups in the PCR products was calculated using the housekeeping gene, *rpl31.* The choice for reference gene was based on preliminary results in the lab where gene expression levels were assessed in embryos overexpressing *dyrk1a* at the same stage (manuscript in prep). The ribosomal gene *rpl31*, was unchanged by *dyrk1a* (other common reference genes such as *gapdh* were altered) and ribosomal genes are often used as reference genes in expression analyses.

## Results

### 1. Patients with DYRK1A loss of function gene variants have craniofacial differences

DYRK1A sequence variants have been associated with DYRK1A Haploinsufficiency Syndrome which is characterized by craniofacial differences. To further demonstrate that DYRK1A could be required for craniofacial development in humans then we analyzed craniofacial phenotypes in patients with other sequence or copy number variants (that are predicted to cause DYRK1A loss of function). DYRK1A is predicted to be intolerant to loss of function and a high probability of being haploinsufficient (gnomAD, pI=1.0, pHaplo=1.0). Despite this, there were 68 patients with loss of function DYRK1A gene variants and their phenotypes described in the Decipher Database. Of the total number of phenotypes reported in patients with DYRK1A loss of function sequence variants, 31.25% were structural changes in the craniofacial region (Fig. 1A). Strikingly, of all the patients identified with loss of function DYRK1A sequence variants, 62.5% had some type of craniofacial phenotype (Fig. 1B). We similarly examined patients with CNVs likely causing loss of function of DYRK1A. Of the total number of phenotypes reported in patients with DYRK1A CNVs, 32.3% were structural changes in the craniofacial region (Fig. 1C). Of all the patients identified with loss of function DYRK1A CNVs (and phenotypes reported), 64.3% had some type of craniofacial phenotype (Fig. 1D). These results are consistent with Dyrk1a Syndrome and support our hypothesis that DYRK1A is required for craniofacial development.

**Figure 1:**
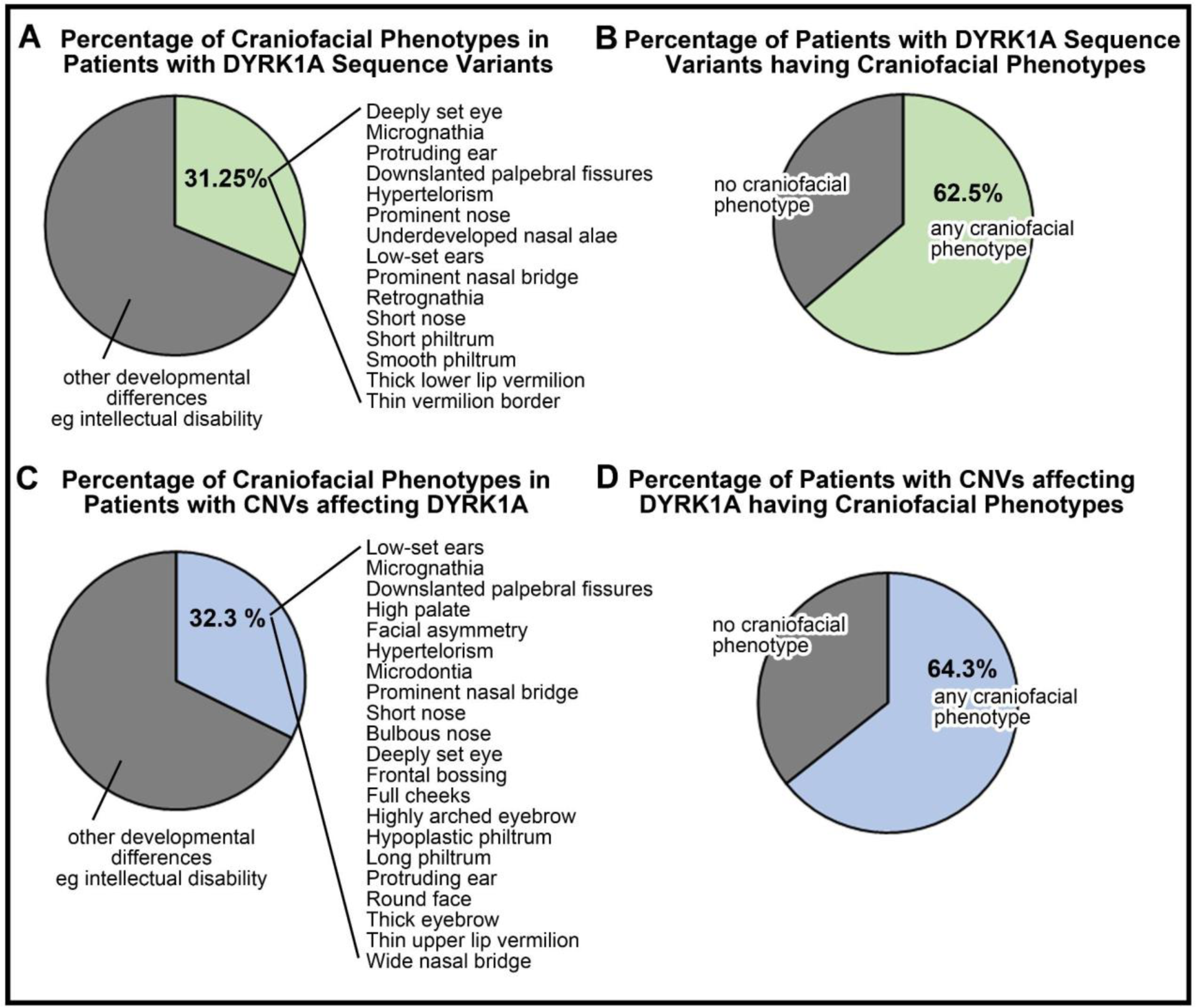
DECIPHER analysis of phenotypes in patients with DYRK1A gene variants. **A)** Pie chart showing that 31.25 % (green) loss of function sequence variants had structural changes in the craniofacial region (n=15/48 patients). **B)** 62.5% (green) of patients identified with loss of function DYRK1A sequence variants had a craniofacial difference (n=25/40 patients). **C)** 32.3 % (blue) of patients with DYRK1A CNVs had structural changes in the craniofacial region (n=21/65). **D)** 64.3% (blue) of patients identified with loss of function DYRK1A CNVs had some type of craniofacial phenotype (n=18/28).

### 2. The X. laevis Dyrk1a proteins are highly similar to the human protein

To test the role of Dyrk1a in craniofacial development we turned to a well established developmental animal model, *Xenopus laevis*. This species is allotetraploid and has two copies of each gene denoted Long (L) and Short (S). Using NCBI Blast we determined that Dyrk1a.L and Dyrk1a.S proteins are 96% similar to each other and are expressed at similar levels over time (suppl Fig. 1). Further, both of these proteins share 94% similarity with the human DYRK1A protein (isoform 2, NP_001334650.1) suggesting that the *X. laevis* proteins are highly conserved (suppl. Fig. 1) and likely to share common functions with the human protein.

### 3. Dyrk1a is expressed in the developing craniofacial region of Xenopus

If Dyrk1a is required for craniofacial development then we expect that *dyrk1a* mRNA and protein would be expressed in developing craniofacial tissues. Indeed, previous work demonstrated that *dyrk1a* transcripts are expressed in the craniofacial region throughout early development (Blackburn et al., 2019; Willsey et al., 2020). Here we have also demonstrated that *dyrk1a* mRNA was expressed in the head, specifically focusing on the early developing face. At stage 25 (∼27hpf) when the neural crest has migrated into the future face, *dyrk1a* was observed throughout the head and slightly enriched in the developing eye (Fig. 2A i,ii). At stage 32 (∼40hpf), the branchial arches, which are reiterated compartmental structures will form the hard and soft tissues of the head including the jaws are present. *dyrk1a* was enriched in these branchial arches as well as the developing facial tissues surrounding the mouth (Fig. 2A iii,iv). At this time, *dyrk1a* also appears to be enriched in the otic vesicle and the brain. At stage 35 (∼50 hpf), *dyrk1a* is enriched in facial tissues surrounding the eye and the mouth that will form the primary palate and anterior most jaw cartilages (Fig. 2A v,vi). Sense controls did not result in staining (suppl. Fig. 2)

**Figure 2:**
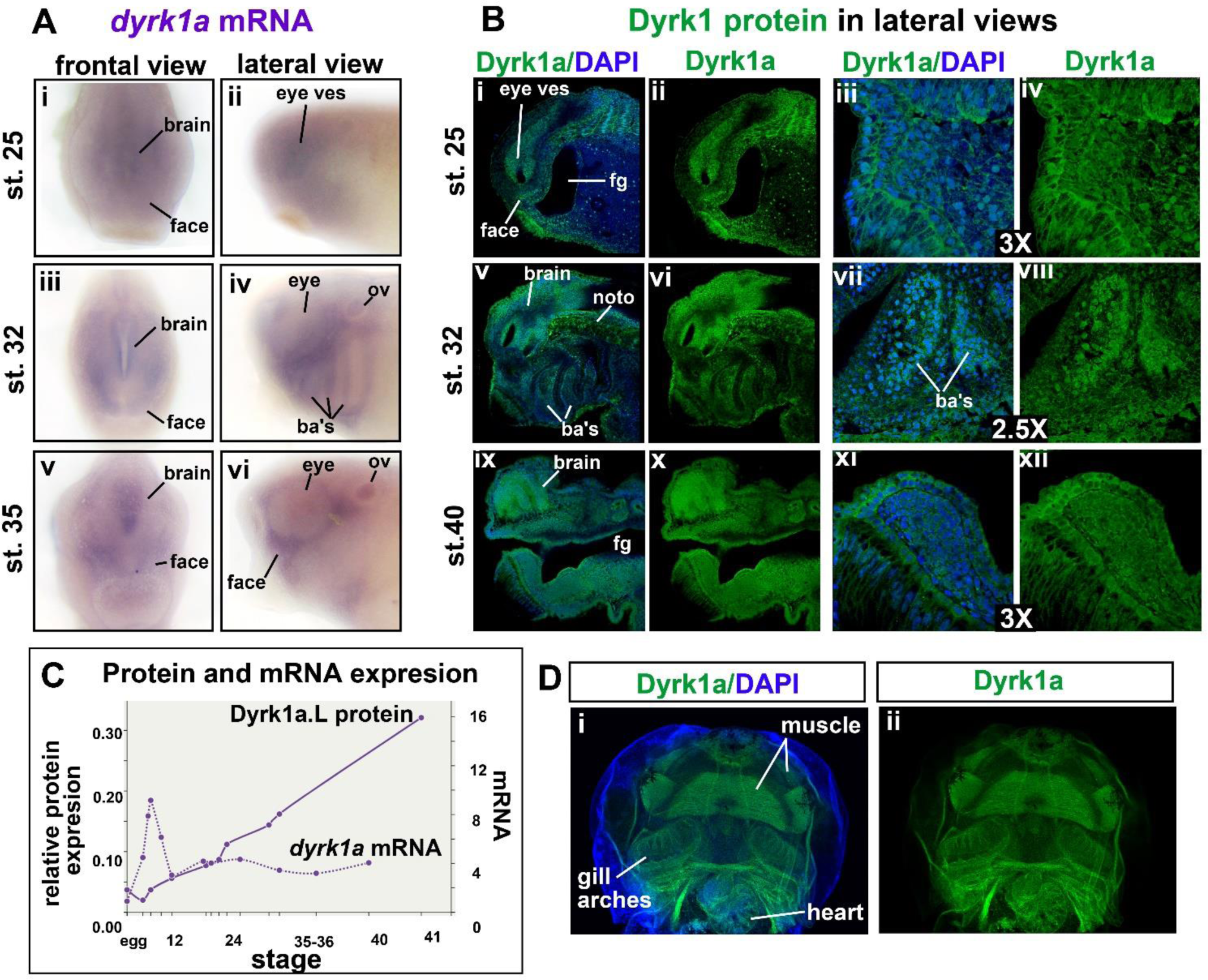
Dyrk1a expression in the craniofacial regions of *X. laevis* embryos. A) Representative frontal and lateral views (anterior to left) of embryos showing *dyrk1* mRNA expression using in situ hybridization at three time points. **i,ii)** stage 25 (n=27, 2 experiments). **iii, iv)** stage 32 (n=29, 2 experiments). **v,vi)** stage 35 (n=30, 2 experiments). **B)** Representative embryos showing Dyrk1 protein expression by immunofluorescence. All images are sagittal sections (anterior to the left) with Dyrk1a labeled green and DAPI labeled blue. Representatives are shown from 18-20 embryos in 2 experiments. **i-iv)** stage 25. **v-viii)** stage 32. **ix-xii)** stage 40. **C)** Plot of relative expression of *dyrk1a.L* mRNA and Dyrk1a.L protein generated in Xenbase.org ((http://www.xenbase.org/, RRID:SCR_003280). D) Ventral view of a representative embryo labeled with Dyrk1a antibody at stage 43 (anterior is to the top, Dyrk1a=green and DAPI=blue). Based on 20 embryos in 2 biological replicates. Abbreviations: ves=vesicle, ba’s=branchial arches, fg=foregut, noto=notochord, ov=otic vesicle.

To visualize Dyrk1a protein expression in Xenopus, we used an antibody that recognizes the C-terminus of the human DYRK1A protein. This region is highly similar to the corresponding region of the *X. laevis* protein sequence (suppl. Fig. 1). When embryos were injected with Xenopus *dyrk1a* mRNA into one cell at the 2-cell stage we noted an increase in DYRK1A antibody labeling in the injected half of the embryo (suppl. Fig 2). Further, this antibody also labeled cilia as other DYRK1A antibodies described in previous reports (Willsey et al., 2020) and suppl. Fig. 2). Together, our data indicates that the antibody specifically labeled the *Xenopus* Dyrk1a protein. At early stages (∼st. 25, 27hpf) we observed Dyrk1a throughout the tissues comprising the head (Fig. 2B i,ii). Enrichment of the protein was observed in the developing eye consistent with the *in situ* hybridization data (Fig. 2B i,ii). Higher magnified images revealed more details about Dyrk1a localization. For example, Dyrk1a appeared to be associated with the plasma membrane as well as nuclei in the epidermal cells (Fig. 2B iii,iv). Later, at stage 32 (∼40hpf), we noted again Dyrk1a was present in a majority of the cells comprising the head (Fig. 2B v,vi). In particular, we observed an enrichment in the mesenchymal core of the branchial arches composed of neural crest cells (Fig. 2B v-viii). Interestingly, in addition to nuclei, the lamellipodia-like protrusions in these cells were strongly labeled with the DYRK1A antibody (Fig. 2B vii-viii). At stage 39-40, as the embryonic mouth is forming, the antibody labeled both the cytoplasm and nuclei of many cells in developing facial tissues (Fig. 2B ix-xii). Finally, Dyrk1a appeared enriched in the craniofacial tissues in later stages (st. 43-44). Specifically, the muscles (also see section 6) as well as the gill arches appeared enriched in Dyrk1a protein (Fig. 2Di,ii).

Using the Xenbase.org database, we plotted *dyrk1a* mRNA and protein levels over time. These results showed relatively constant levels of *dyrk1a* mRNA after gastrulation when critical events in craniofacial take place (Fig. 2C). On the other hand, Dyrk1a protein steadily increased over the same period of time (Fig. 2C). The disconnect in the pattern of mRNA and protein levels could be caused by various factors and additional experiments would be necessary to elucidate these. However, it does emphasize the importance of examining protein expression in addition to mRNA during development.

Together, this expression data shows that Dyrk1a is present throughout the developing craniofacial tissues in both nuclear and cytoplasmic compartments suggesting that it could have multiple roles in multiple tissues in the developing face.

### 4. Decreased Dyrk1a results in craniofacial malformations in Xenopus

To definitively show that Dyrk1a is required for craniofacial development, we next aimed to decrease its function and assess craniofacial morphology. To accomplish this, we used validated antisense oligos (Morpholinos; MOs) and pharmacological tools that have been shown effective in *Xenopus* embryos.

The *dyrk1a* or standard control MOs (Genetools) were injected into the one cell stage and the faces examined at stage 43 (Fig. 3Ai). We noted increased severity of craniofacial malformations with increasing concentration of the MOs. At the lowest dose of dyrk1a MO (5ng) the faces appeared smaller and mouth shape was more oval (Fig. 3A ii,iii). At 10 ng of MO per embryo, the eyes were closer together, the midfaces appeared narrower, and the mouths were rounder (Fig. 3A iv). At 20 ng per embryo, the faces were very narrow, so much so that the eyes appeared fused and the mouth was very small (Fig. 3A v).

**Figure 3:**
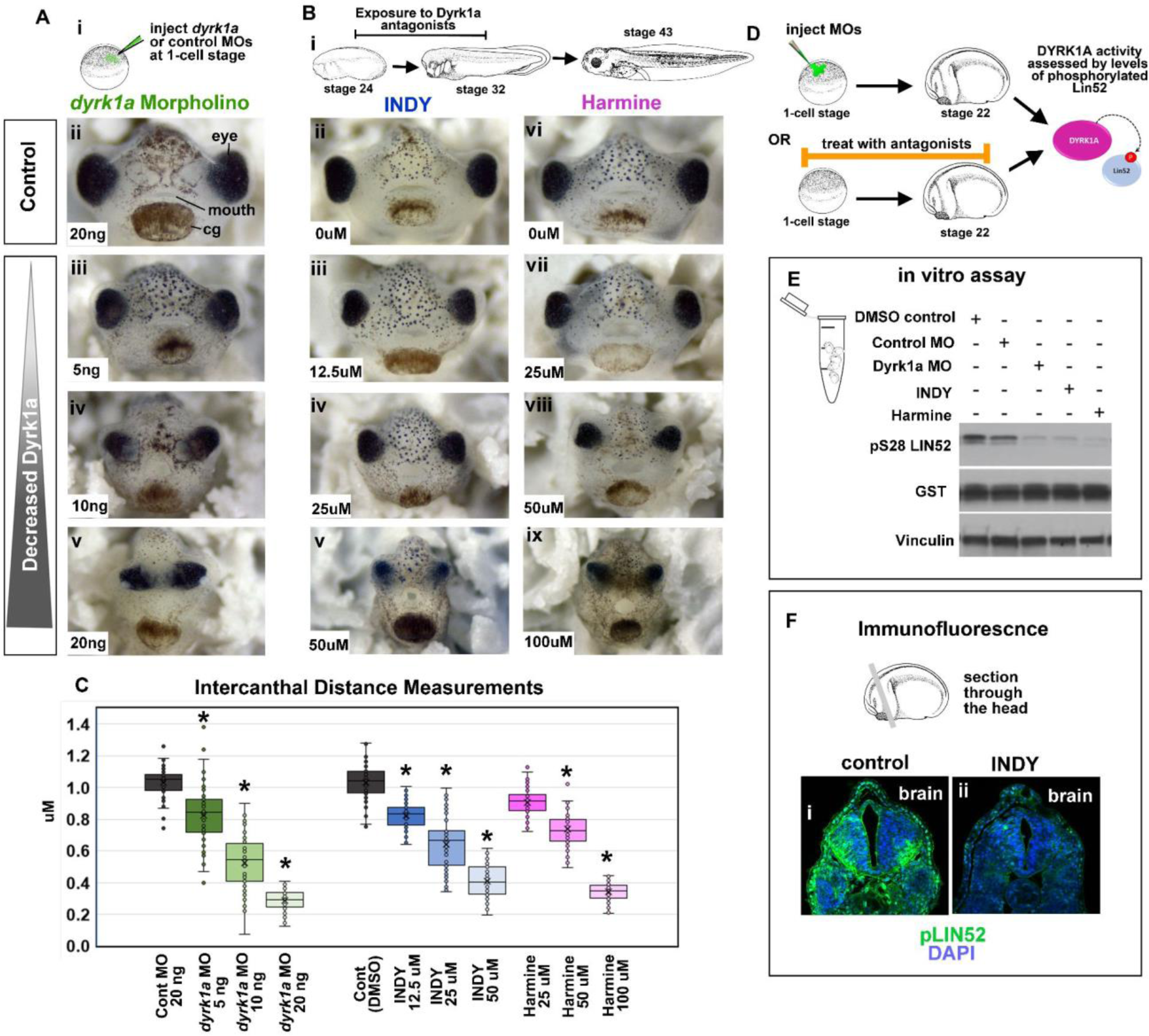
Decreased Dyrk1a function causes craniofacial differences. **Ai)** Schematic showing *dyrk1a* MO injection. Aii-v) Frontal views of representative embryos injected with increasing concentrations of *dyrk1a* MOs. **Bi)** Schematic showing Dyrk1a inhibitor treatments. **Bii-ix)** Frontal views of representative embryos treated with increasing concentrations of INDY or harmine. **C)** Quantification of the intercanthal distance (distance between the eyes) based on 60 embryos in 3 biological replicates (ANOVA, TUKEY posthoc tests, all p values <0.001). **C)** Schematic showing DYRK1A in vitro activity assay protocol to assess effectiveness of Dyrk1a knockdown tools. **E)** Western blots showing phosphorylated LIN52 on serine residue 28. Vinculin was used as a loading control. **Fi-ii**) Representative tissue sections through the head in control (i) and INDY treated embryos (ii). Based on 10 embryos in 2 biological replicates.

To help demonstrate that the effects of the *dyrk1a* MO on craniofacial development were specific to loss of Dyrk1a function we next tested two Dyrk1a inhibitors, INDY and harmine. Embryos were treated from stage 24 until stage 32 (Fig. 3B i) which provides the extra benefit of informing whether the *dyrk1a* MO induced craniofacial malformations are due to earlier effects on gastrulation or neural crest specification. Results indicated that the Dyrk1a inhibitors had very similar effects as the *dyrk1a* morpholinos on craniofacial development. Embryos had narrower faces, closer set eyes and rounder mouths. The severity of these defects also increased with increasing concentrations (Fig 3B ii-ix). Since decreased Dyrk1a caused narrower appearing faces we measured intercanthal distance (the distance between the eyes) in our morpholino and inhibitor experiments. This quantification closely reflected our observations, where *dyrk1a* MOs as well as INDY and harmine treatments caused a decrease in the intercanthal distance that was more profound as concentrations increased (Fig. 3C).

### 5. Tools to decrease Dyrk1a in Xenopus prevent phosphorylation of a Dyrk1a target LIN52

It is possible that *dyrk1a* MOs, INDY and harmine cause a phenotype through one or more non-specific mechanisms rather than decreasing Dyrk1a function. Therefore, we next validated that these methods did indeed knockdown Dyrk1a function in the embryo. To do so we used an established in vitro activity assay that assessed the phosphorylation of a verified DYRK1A target, LIN52 (Iness et al., 2019; Menon et al., 2019) (Fig 3D). Results indicated that *dyrk1a* MOs as well as embryos treated with Dyrk1a inhibitors reduced the phosphorylation of LIN52 in embryos (Fig. 3E). Using immunofluorescence we also noted a decrease in phosphorylated Lin52 in the head of embryos after INDY exposure (Fig 3F i,ii). Together these results suggested that our tools are indeed effective at reducing Dyrk1a function.

### 6. Decreased Dyrk1a results in abnormalities in cranial cartilages and muscles

The jaw cartilage and muscle provide the shape of the face and therefore we examined these tissues in embryos with decreased Dyrk1a function. Embryos were injected with *dyrk1a* MO or exposed to the Dyrk1a inhibitor, INDY (Fig. 4A,B). Cartilages, labeled with Alcian blue staining, were dramatically reduced in 100% of both embryos injected with *dyrk1a* MOs or treated with INDY (Fig. 4C i-iii). Similarly, jaw muscles labeled with a muscle specific antibody (12/101) were also reduced in 100% of the embryos examined (Fig. 4C iv-vii). To determine whether Dyrk1a might have a direct role in cartilage and muscle development we co-labeled Dyrk1a with either collagen II or muscle (12/101). In tissue sections labeled with a collagen II antibody we noticed positive staining surrounding cartilage elements. Dyrk1a labeling overlapped with collagen II and appeared within the cartilage elements as well. Specifically, Dyrk1a appeared enriched in the nuclei of cells inside of the cartilages (Fig. 4D i-iii) suggesting a nuclear role in these tissues. Muscle specific antibody,12/101, labels a membrane protein of the sarcoplasmic reticulum and therefore appears in the cytoplasmic compartment of muscle cells. Dyrk1a appeared to overlap with 12/101 in the muscle (Fig. 4E i-iii) suggesting a cytoplasmic role in this tissue. Future experiments are required to determine the role of Dyrk1a in these tissues.

**Figure 4:**
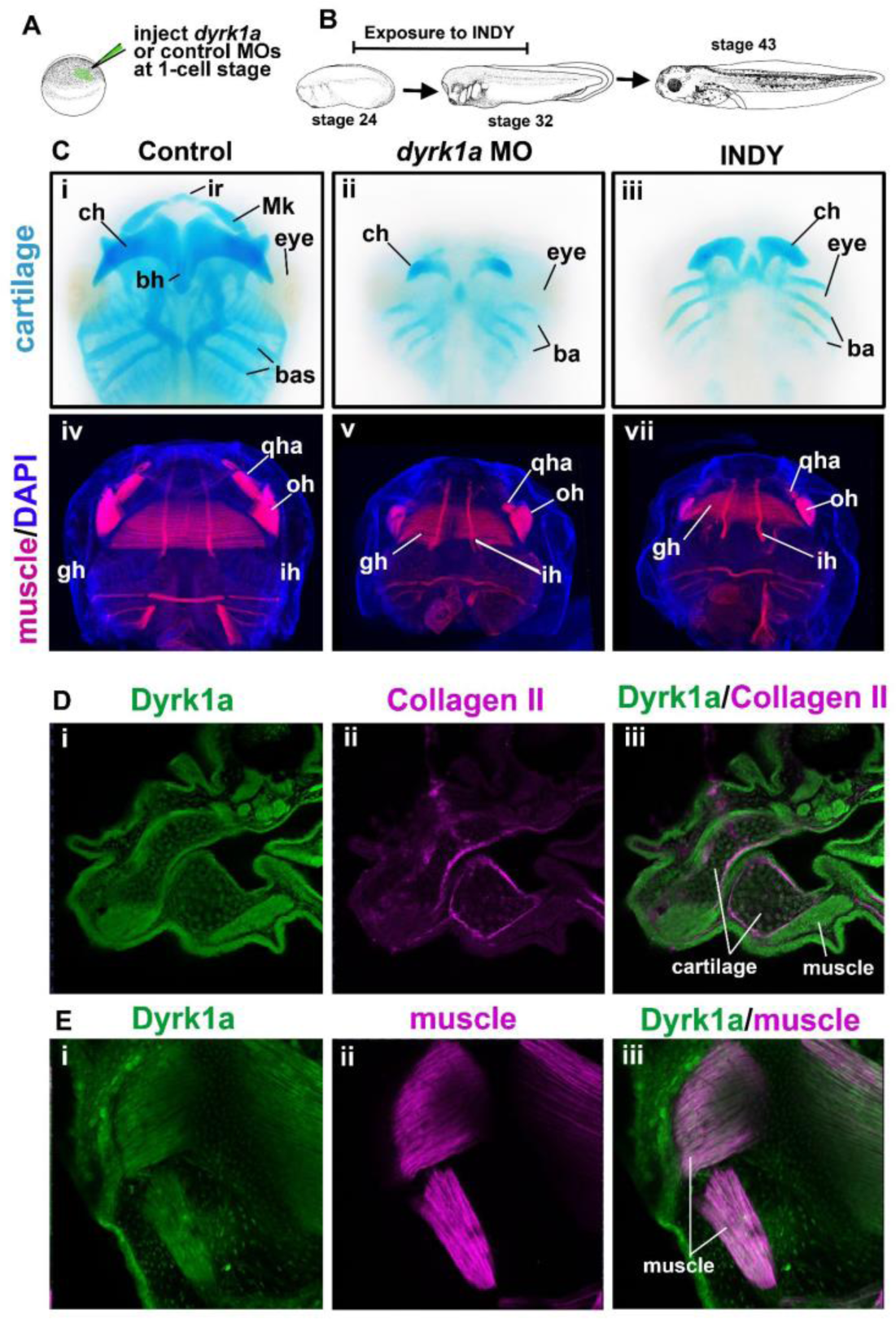
Dyrk1a in jaw cartilage and muscle development. **A)** Schematic of the injection protocol. **B)** Schematic of the exposure protocol. **Ci-iii)** Alcian blue labeling of cartilages of representative embryos (ventral views) injected with dyrk1a MOs or treated with INDY. Based on examination of 40 embryos (2 biological replicates). Abbreviations: cg=cement gland, ch=ceratohyal, ir= infrarostral, Mk=Meckel’s, br=branchial. **Civ-vii)** Muscle specific labeling (12/101) of representative embryos (ventral views) injected with dyrk1a MOs or treated with INDY. Based on examination of 36 embryos (2 biological replicates). Abbreviations; qha= quadratohyoangularis, oh= orbitohyoideus, lm=levator mandibulae, gh= geniohyoidues. **Di-iii)** Double labeling of a representative tissue section through the head. Dyrk1a is labeled green and Collagen II is labeled pink. Collagen II labels cells of the craniofacial cartilages. Based on 20 embryos (2 biological replicates). **Ei-iii)** Double labeling of a representative tissue whole mount embryo of the head. Dyrk1a is labeled green and muscle (12/101) is labeled pink. Based on 20 embryos (2 biological replicates)

### 7. Decreased Dyrk1a function alters expression of craniofacial regulators and neural crest

Craniofacial development is regulated by a number of signals and transcription factors. We next assessed whether the expression of craniofacial regulators were altered in response to decreased Dyrk1a function. We chose to focus on a small subset of genes that overlapped with dyrk1a mRNA expression in the branchial arches or head at stages 30-35 (Xenbase.org) and represented transcription factors and retinoic acid signaling. The heads of embryos exposed to INDY were isolated, RNA extracted and RT-qPCR performed (Fig. 5A). Results indicated that there were reductions in *sox9* (0.61 fold), *pax3* (0.38 fold), *alx4* (0.65 fold) and *rarg* (0.49 fold) and a minor reduction in *gbx2* (0.82 fold) in the head tissues of embryos exposed to INDY relative to control untreated embryos (Fig. 5B). On the other hand there was increased expression of *crabp2* (2.5 fold) in these same tissues (Fig. 5B). These results indicate that Dyrk1a is required for the proper expression of specific craniofacial regulators expressed in the branchial arches. Additionally, this data also demonstrates that decreased Dyrk1a doesn’t simply cause a global reduction in gene expression that might be caused by sick or unhealthy embryos.

**Figure 5:**
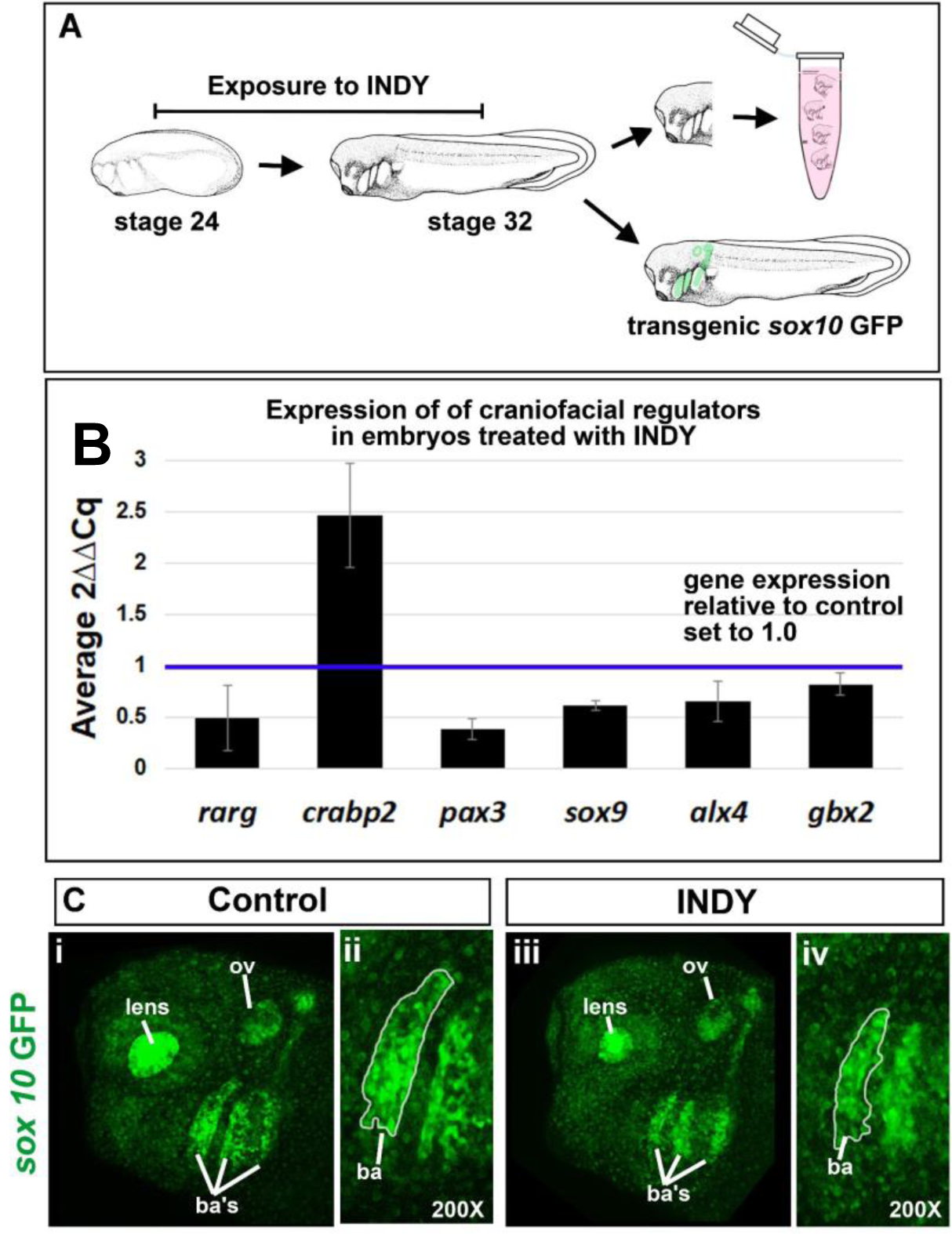
Decreased Dyrk1a function results in changes to craniofacial regulators and neural crest. **A)** Schematic of the experimental plan. **B)** Results of RT-qPCR showing expression levels of craniofacial regulators. Levels are shown relative to controls set to 1 (blue line). Two biological replicates were performed and variation between the two is represented by the error bar (standard error). Each biological replicate is an average of 3 technical replicates. Error bars are *alx4* = 0.20, *crabp2* = 0.50, *gbx2* = 0.11, *pax3* = 0.11, *rarg* = 0.32, and *sox9* =0.05. C) i) Lateral view of the head, anterior to the left, of a representative *sox10*-GFP transgenic embryo exposed to control DMSO. ii) Magnified image of the anterior branchial arch sox10 expression domain in i outlined in white. iii) Lateral view of the head, anterior to the left, of a representative *sox10*-GFP transgenic embryo exposed to INDY. iv) Magnified image of the anterior branchial arch *sox10* expression domain in iii outlined in white.

Both *pax3* and *sox9* are important for cranial neural crest development. Therefore, since the expression levels of these genes were dramatically reduced we next assessed the neural crest visually. To do this, a transgenic line where GFP is expressed under the *sox10* promoter (a gene expressed in cranial neural crest) was assessed (Alkobtawi et al., 2018; Ossipova et al., 2018). These transgenic embryos were exposed to INDY from stage 24 to 32 and the GFP examined (Fig. 5A). The results indicated that the GFP expression pattern appeared very similar in control and INDY treated embryos (Fig. 5C i-iv). However, the size of each the *sox10* expression domain in the branchial arches appeared smaller in INDY treatments compared to the controls (Fig. 5C i-iv). It is possible that Dyrk1a is required for the correct numbers of cranial neural crest cells in the branchial arches.

### 8. Decreased Dyrk1a function results in increased cell death and decreased proliferation in the developing face

Dyrk1a is known to regulate numerous processes including cell cycle and apoptosis. Therefore, we next wondered whether craniofacial defects were due to an excess in cell death and/or a lack of proliferation in the tissues that form the face.

To test whether decreased Dyrk1a altered levels of apoptosis we used a well characterized marker, cleaved caspase-3 (CC3). *dyrk1a* MOs (or control MOs) were injected into one cell at the 2-cell stage to isolate MOs to one half of the embryo. This was followed by CC3 labeling at stage 26-27, a time before MOs are highly diluted (Fig. 6A). CC3 labeling at the resolution detected was not amenable to counting individual cells. Therefore, we calculated CC3 positive fluorescence of the head, excluding the eyes and brain. The fluorescence intensity was then normalized to the area and values plotted. Results indicated that there was no significant difference in CC3 labeling on the control morphant side of the head compared to the uninjected side of the head (Fig. 6B i-iv). On the other hand, we did determine that CC3 induced fluorescence was significantly higher on the *dyrk1a* morphant side of the head compared to the control side of the head (Fig. 6B v-viii).

**Figure 6:**
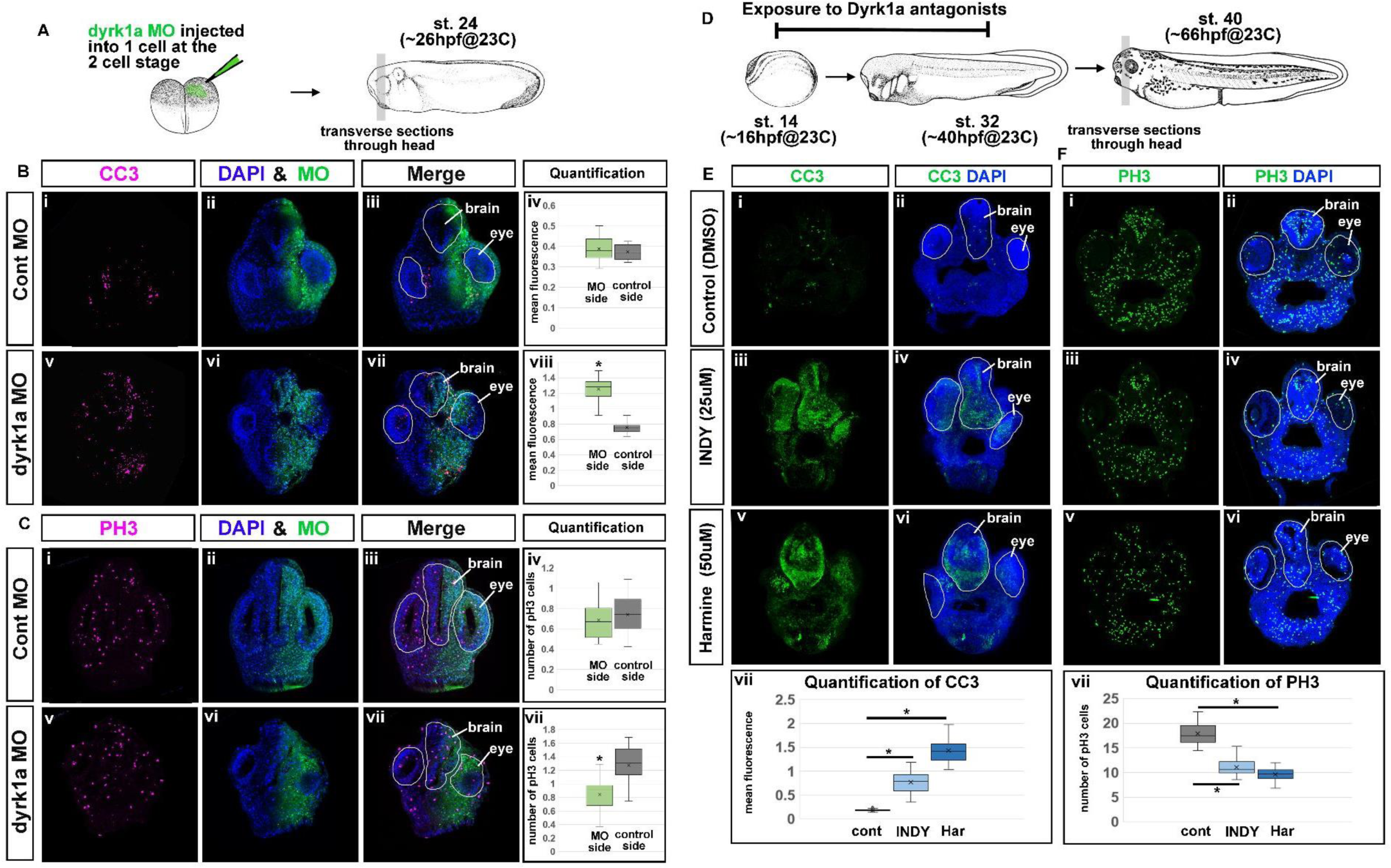
Decreased Dyrk1a function alters cell survival and proliferation. **A)** Schematic shows the experimental design where fluorescein labeled MOs (dyrk1a or control) were injected in one cell at the 2-cell stage and then embryos were fixed, sectioned and labeled. **Bi-iii)** Representative sections showing CC3 labeling (pink), Control MO (green) and DAPI counterstain. **Biv)** Quantification of 20 embryos (2 replicates) showing no statistical difference between CC3 labeling on the control MO injected sides versus the uninjected sides (ttest, p=0.314). **Bv-vii)** Representative sections showing CC3 labeling (pink), *dyrk1a* MO (green) and DAPI counterstain. **Bviii)** Quantification of 20 embryos (2 replicates) showing a statistical difference between labeling on the *dyrk1a* MO injected sides verses the uninjected sides (ttest, *p<0.001). **Ci-iii)** Representative sections showing PH3 labeling (pink), Control MO (green) and DAPI counterstain (blue). **Civ)** Quantification of 20 embryos (2 replicates) showing no statistical difference between CC3 labeling on the control MO injected sides versus the uninjected sides (ttest, p=0.333). **Cv-vii)** Representative sections showing PH3 labeling (pink), *dyrk1a* MO (green) and DAPI counterstain (blue). **Cviii)** Quantification of 20 embryos (2 replicates) showing a statistical difference between labeling on the *dyrk1a* MO injected sides verses the uninjected sides (ttest, *p<0.001). **D)** Schematic of the experimental design where embryos were treated with INDY or Harmine and fixed, sectioned and labeled. **Ei-vi)** Representative sections from embryos, treated with INDY or Harmine, showing CC3 labeling (green), and DAPI counterstain CC3 (green). **Evii)** Quantification of 20 embryos (2 replicates) shows statistical differences in CC3 labeling between control and INDY and control and Harmine (Kruskal-Wallis One Way Analysis of Variance on Ranks p<0.001, TUKEY posthoc tests, control vs INDY *=p<0.001 and control vs harmine *=p<0.001). **Fi-vi)** Representative sections from embryos, treated with INDY or Harmine, showing PH3 labeling (green), and DAPI counterstain. **Fvii)** Quantification of 20 embryos (2 replicates) shows statistical differences in PH3 labeling between control and INDY and control and Harmine (Kruskal-Wallis One Way Analysis of Variance on Ranks p<0.001, TUKEY posthoc tests, control vs INDY *=p<0.001 and control vs harmine *=p<0.001).

To test whether decreased Dyrk1a altered proliferation we used a well characterized marker of cell division, phosphorylated histone H3 (PH3). *dyrk1a* MOs (or control MOs) were injected into one cell at the 2-cell stage as described above (Fig. 6A). This was followed by PH3 labeling, also at stage 26-27. PH3 labeling allowed us to count individual cells and therefore we calculated cells in the head, excluding the eyes and brain which was then normalized to the area. Results indicated that there was no significant difference in PH3 positive cells on the control morphant side of the head compared to the uninjected side of the head (Fig. 6Ci-iv). Conversely, we determined that the number of PH3 positive cells was significantly lower on the *dyrk1a* morphant side of the head compared to the control side of the head (Fig. 6C v-viii).

While not as common as in zebrafish, morpholinos can sometimes cause non-specific cell death. Therefore, we asked whether there is a similar increase in CC3 labeling and conversely a decrease in PH3 positive cells in embryos treated with Dyrk1a inhibitors. We also used this opportunity to test whether these effects also occurred over a specific time window when the facial structures are forming after gastrulation is completed (Fig. 6D). Indeed, we determined that there was a significant increase in CC3 positive fluorescence in embryos exposed to INDY or Harmine during early craniofacial development (Fig. 6E i-vii). Additionally, also consistent with the results in *dyrk1a* morphants, PH3 positive cells were also decreased in embryos exposed to both INDY and Harmine (Fig. 6F i-vii). Together, these results indicate that decreased Dyrk1a function could cause a reduction in cell numbers in the face which could account for the narrowing of the faces, reduced cranial neural crest cells, and smaller jaw cartilages and muscle.

## Discussion

Craniofacial development is a complex orchestration of signaling, growth and differentiation. It is therefore no surprise that Dyrk1a, a ubiquitous kinase regulating a wide range of basic cellular processes, would have a role in facial formation. Certainly, we and others have shown that Dyrk1a is expressed in the craniofacial region throughout development in *Xenopus* and mice (Blackburn et al., 2019; Fotaki et al., 2002; Willsey et al., 2020). Moreover, here we demonstrate that embryos, with reduced Dyrk1a, have craniofacial malformations early in development.

Variants in DYRK1A, resulting in haploinsufficiency of the gene, are associated with DYRK1A Syndrome. This syndrome is defined by intellectual disability and autism related disorder, however patients also present with a number of craniofacial differences (Ji et al., 2015; van Bon BWM, 2015). Further analysis of phenotypes in people with various DYRK1A sequence and copy number variants indicate that in fact the majority of such patients have craniofacial differences. In an animal model we have now demonstrated that decreasing Dyrk1a function in *Xenopus* embryos results in malformations consistent with the craniofacial differences observed in humans with such Dyrk1a variants. For example, humans with DYRK1A syndrome often have a smaller head size (microcephaly), narrower forehead, smaller midface features (short nose, and philtrum) as well as a smaller chin. In *Xenopus* with deficient Dyrk1a function we also observed a smaller head and midface hypoplasia resulting in a narrower face. Hypoplasia of embryonic tissues have also been noted in other embryos with a reduction in *Dyrk1A* (Fotaki et al., 2002; Kim et al., 2017)(Blackburn et al., 2019; Fotaki et al., 2002; Willsey et al., 2020). Together, these and our own data suggest that without Dyrk1a function, embryos have a deficiency in the size of embryonic organs. In *Xenopus*, we have begun to determine why there is a deficit in the size of the face when Dyrk1a function is reduced. First, we know that the shape of the face is largely determined by the size and position of the muscular and skeletal elements. Correspondingly, we determined that embryos with decreased Dyrk1a had both reduced cranial cartilage and muscle. Dyrk1a may have a direct role in the development of these structures since we determined that this protein is present in cartilage and muscle cells. Certainly, DYRK1A has been shown to have a role in myogenesis further supporting this possibility in muscle (Yu et al., 2019). Dyrk1a may also affect the development of jaw muscle and cartilage indirectly by modulating processes important for their development such as gene expression as well as growth and cell survival.

In terms of gene expression, embryos with reduced Dyrk1a function did indeed have altered expression of critical craniofacial regulators. Transcription factors *pax3* and *sox9* were decreased in the facial tissues of embryos treated with a Dyrk1a inhibitor. Mutations in *PAX3* cause Craniofacial-deafness-hand syndrome (MIM 122880) characterized by craniofacial differences including short nose, short palpebral fissures and hypertelorism (Asher et al., 1996). Additionally, *SOX9* mutations cause Campomelic dysplasia (MIM 114290), which causes defects in skeletogenesis, but patients also have craniofacial differences such as midface hypoplasia. Both *pax3* and *sox9* are required for cranial neural crest development, an important component of the developing face (Lee and Saint-Jeannet, 2011; Maczkowiak et al., 2010; Monsoro-Burq et al., 2005; Spokony et al., 2002). A transcription factor, *alx4*, required for the development of the region around the nose and forehead was also reduced in the heads of embryos deficient in Dyrk1a function (Beverdam et al., 2001; Hussain et al., 2020). In addition, two members of the retinoic acid signaling pathway were also altered in embryos with decreased Dyrk1a function. While there was a reduction in the retinoic acid receptor, *rarg*, there was an increase in the retinoic acid binding protein *crabp2*. Dyrk1a could be a negative regulator *crabp2* or alternatively, upregulation of this gene could be a compensation mechanism in response to the depletion of *rarg* in the developing face. Previous work has shown that retinoic acid signaling is required for orofacial development across vertebrates (Dubey et al., 2018; Morriss-Kay, 1993; Williams and Bohnsack, 2019; Wu et al., 2022). In *Xenopus*, reductions in this pathway can cause orofacial clefts, slanted eyes and midface hypoplasia and embryos deficient in Dyrk1a function also have similar malformations (Kennedy and Dickinson, 2012, 2014). In summary, misregulation of any of the investigated fundamental craniofacial regulators could perturb growth and differentiation of the facial tissues and account for craniofacial malformations observed in embryos with deficient Dyrk1a.

Each of the craniofacial regulators altered are expressed in the branchial arches (Karimi et al., 2018) overlapping with *dyrk1a* expression and therefore we hypothesized that dyrk1a could regulate their expression directly. DYRK1A could potentially modulate transcription of the craniofacial regulators by several mechanisms. One possibility is that Dyrk1a phosphorylates upstream signaling proteins required for the expression of these genes. Indeed, DYRK1A phosphorylates proteins integral to several developmental signaling pathways such as hedgehog, nuclear factor of activated T cells (NFAT), Notch, as well as WNT signaling (Arron et al., 2006; Atas-Ozcan et al., 2021; Ehe et al., 2017; Fernandez-Martinez et al., 2009; Hammerle et al., 2011; Hong et al., 2012). However, whether Dyrk1a regulates one or more signaling pathways in the developing face needs further exploration. DYRK1A has also been reported to regulate transcription by its ability to directly phosphorylate RNA polymerase II and histone components and therefore it could also be necessary for the function of transcriptional machinery (Di Vona et al., 2015; Jang et al., 2014; Yu et al., 2019). Our results warrant further experiments to decipher if and how Dyrk1a could be regulating gene expression during craniofacial development.

An important component of craniofacial development is the cranial neural crest. In embryos with decreased Dyrk1a function we noted a significant decrease in genes (*sox9 and pax3*) required for the development of this tissue. We also assessed neural crest in vivo using a transgenic line where GFP expression is driven by another cranial neural crest regulator, *sox10* (Alkobtawi et al., 2018; Gfrerer et al., 2013). Exposure to a Dyrk1a inhibitor did not prevent proper neural crest migration, however, the regions of *sox10* expression appeared smaller. This could be due to fewer neural crest cells in the branchial arches. Thus, it is possible that changes in craniofacial gene expression as well as the smaller size of the face, jaw muscle and cartilage could be explained by reduction in the number of cells during craniofacial development.

One reason for fewer numbers of cells in the developing face could be a decrease in cell survival. Consistently, we found that *Xenopus* embryos with decreased Dyrk1a function do have a dramatic increase in apoptosis in the developing head, indicating that this protein could indeed be required for cell survival during face development. Dyrk1a has also been shown to have a similar role in cell survival in the developing brain and retina (Kim et al., 2017; Laguna et al., 2008; Willsey et al., 2020). Dyrk1a can promote cell survival by inactivating Caspase-9 via phosphorylation of the Thr125 site (Barallobre et al., 2014; Laguna et al., 2008). In doing so, Dyrk1a prevents the activation of the intrinsic apoptotic pathway and prevents inappropriate cell death. Consistently, inactivation of Caspase-9 has been described in human cell lines as a mechanism to promote cell survival (Jeong et al., 2011). Another mechanism by which Dyrk1a could be influencing apoptosis is via its regulation of pro-survival protein SIRT1 (Guo et al., 2010). In human cells, DYRK1A can activate SIRT1 which in turn promotes deacetylation and the activation of p53. As a result, Dyrk1a acts to suppress p53 mediated cell death. Dyrk1a could also influence cell survival indirectly by its role in DNA repair. For example, DYRK1A associates with a DNA repair protein, RNF169, and thereby promotes homologous recombination repair (Menon et al., 2019; Roewenstrunk et al., 2019). In summary, Dyrk1a function could be essential to maintain cell survival by several mechanisms. Future experiments are required to pinpoint the precise means by which cells die in the developing facial tissue when this protein is not present.

DYRK1A could also regulate cell numbers and growth in the face via its roles in the cell cycle. This protein has actually been shown to suppress cell cycle by modulating several cell cycle regulators such as D cyclins, CDK inhibitors, p53, the LIN52 subunit of DREAM complex as well as the spindle microtubules (Fernandez-Martinez et al., 2015; Iness et al., 2019; Litovchick et al., 2011; Liu et al., 2014)(Ananthapadmanabhan et al., 2023). Therefore, Dyrk1a deficiency might then be expected to increase proliferation in the developing embryo. Certainly, this is true in the developing *Xenopus* brain where decreased Dyrk1a results in the increase in cell cycle markers (Willsey et al., 2020). On the other hand, Dyrk1a inhibition has also been shown to suppress cell cycle progression. For example, in glioblastoma cells a moderate level of DYRK1A inhibition leads to increased proliferation as expected, but in contrast, high levels of DYRK1A inhibition results in cell cycle arrest (Recasens et al., 2021). Therefore, how Dyrk1a regulates cell proliferation could depend on the context and dose of this protein. In our experiments we noted a minor but significant decrease in mitotic marker phospho-histone H3 in the faces of embryos injected with *dyrk1a* MO or exposed to Dyrk1a inhibitors. This could be due to a direct inhibitory effect on cell cycle progression or perhaps an indirect effect caused by excess apoptosis and/or decreased expression of craniofacial regulators.

Together, our results indicate that a reduction in Dyrk1a function causes craniofacial defects including a narrower face, smaller jaw cartilage and muscle and decreased cranial neural crest expression domains. These changes correlate with fewer mitotic cells and a significant increase in apoptosis in the developing facial tissues. Our work demonstrates the possibility that embryos with a lower dose of Dyrk1a have an insufficient number of cells which affects the morphology of the face. This insufficiency could be due to a number of mechanisms affecting cell survival, cell proliferation and gene expression and could be further elucidated in future work.

## Acknowledgements

I would like to thank: 1) Nicole Cheng, a lab technician and undergraduate student researcher who took pictures and helped quantify of orofacial morphology. 2) Animal care technicians Deborah Howton and Leona Bhandari. 3) Allyson Kennedy for identifying Dyrk1a in a screen and performing the first INDY exposures in the lab. We are also very thankful to the VCU Children’s Hospital Research Institute for providing funding that allowed us to finish this work.

## Author contributions

**Johnson**: editing, writing and gene expression experiment.

**Wahl**: in situ hybridizations of dyrk1a.

**Sesay:** In vitro DYRK1A activity assay.

**Litovchick:** reviewing, editing and writing.

**Dickinson:** conception, compiled data, writing, reviewing performed experiments in figures 2,3,4,5,6.

---

**Supplemental Figure 1:**
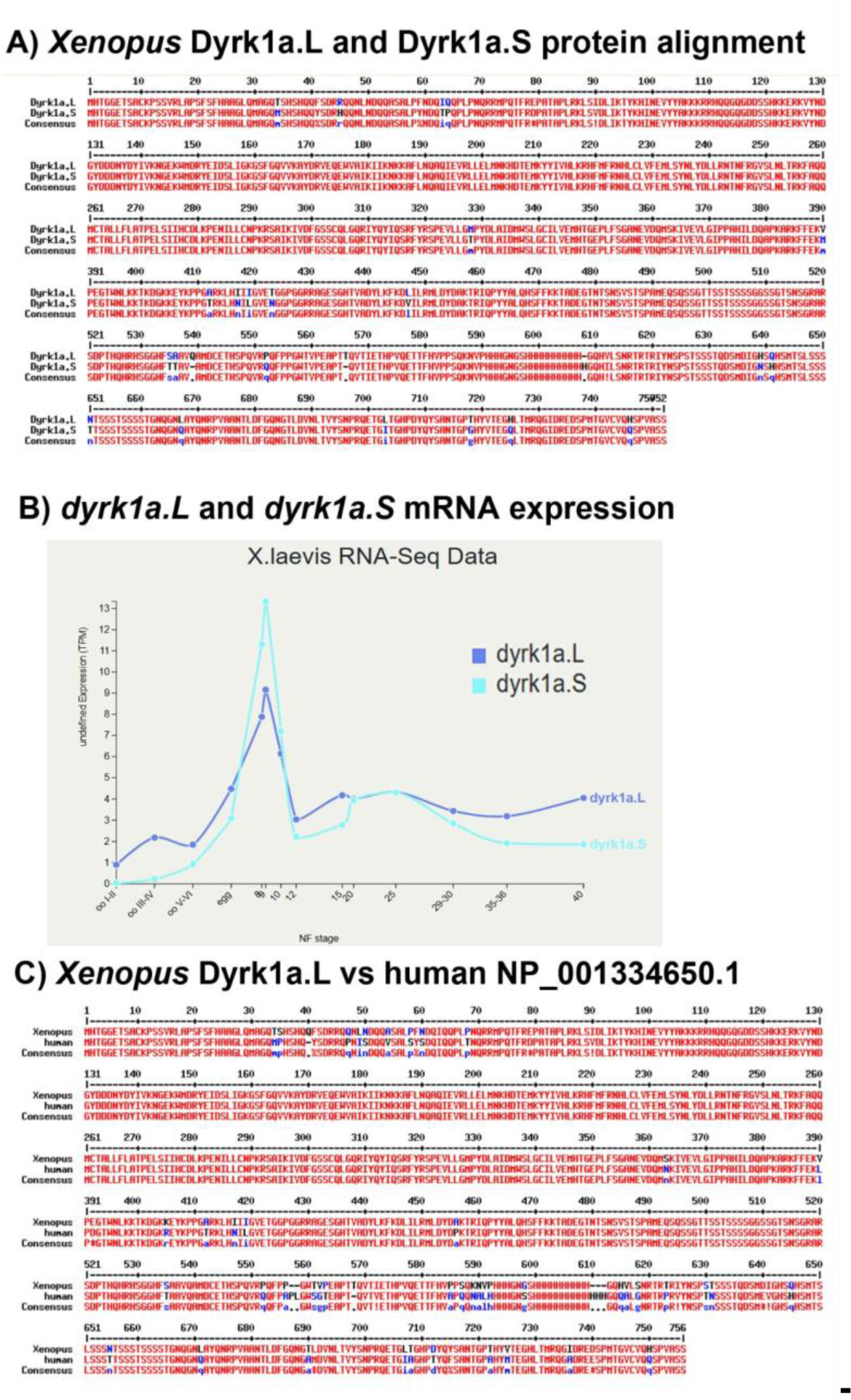
**A)** Dyrk1a.L and Dyrk1a.S protein alignment (96% similarity). **B)** Relative dyrk1a.L and dyrk1a.S mRNA expression over time (generated by Xenbase.org, (Bowes et al., 2008)). **C)** Dyrk1a.L and human DYRK1A protein (isoform 2, NP_001334650.1) alignment (94% similarity).

**Supplemental Figure 2:**
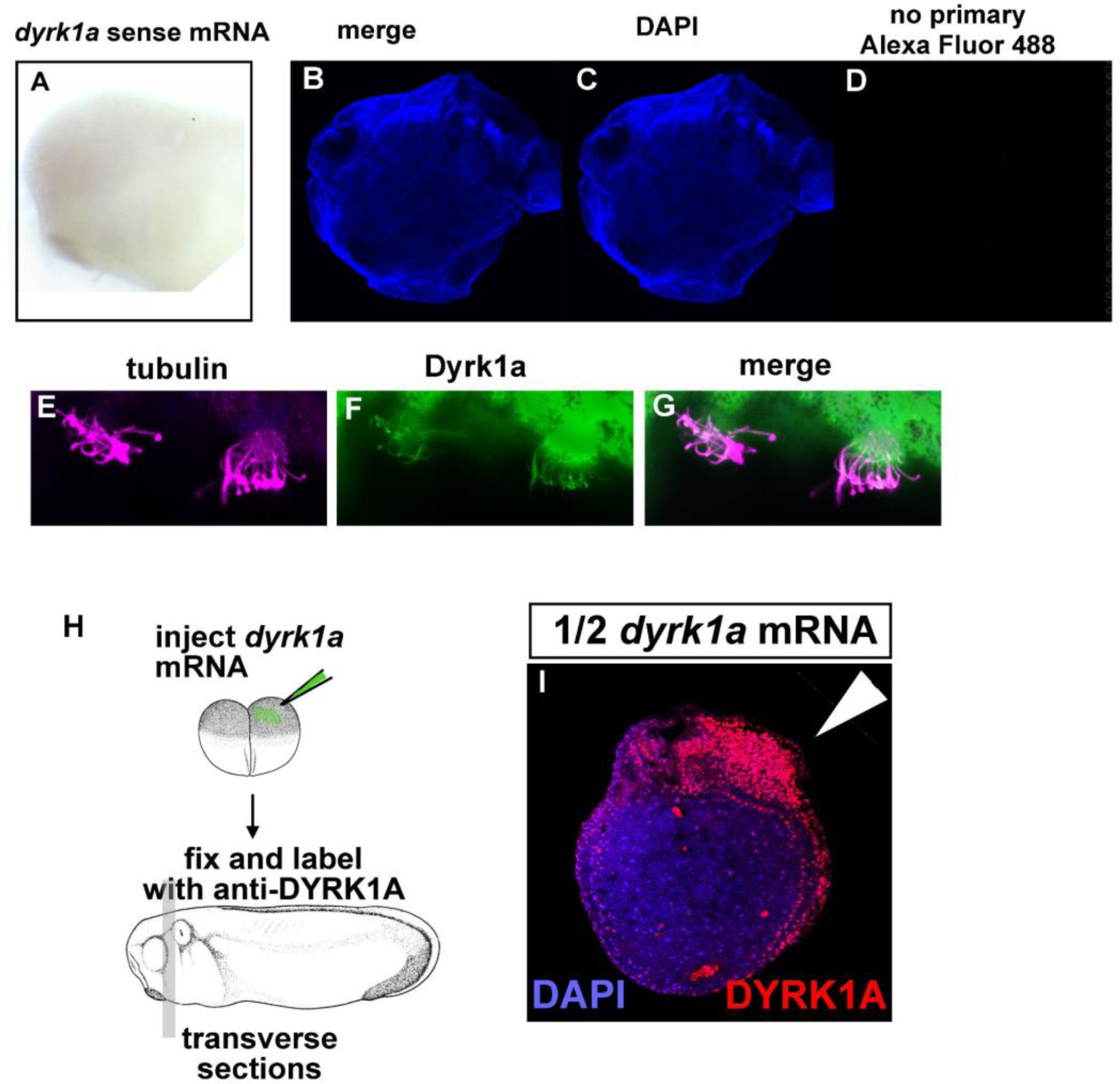
A) Sense control probe for *dyrk1a* does not label tissues. B-D) Immunofluorescence controls where no DYRK1A primary antibody was added shows no signal at the same confocal settings used to collect images where primary was included. E-G) Dyrk1a and alpha tubulin double labeling indicates DYRK1A antibody labels cilia. H) Schematic to show the experimental plan to overexpress dyrk1a in half the embryo. I) DYRK1A antibody labeling (red) on the right side (white arrowhead).

**Supplemental Table 1.**
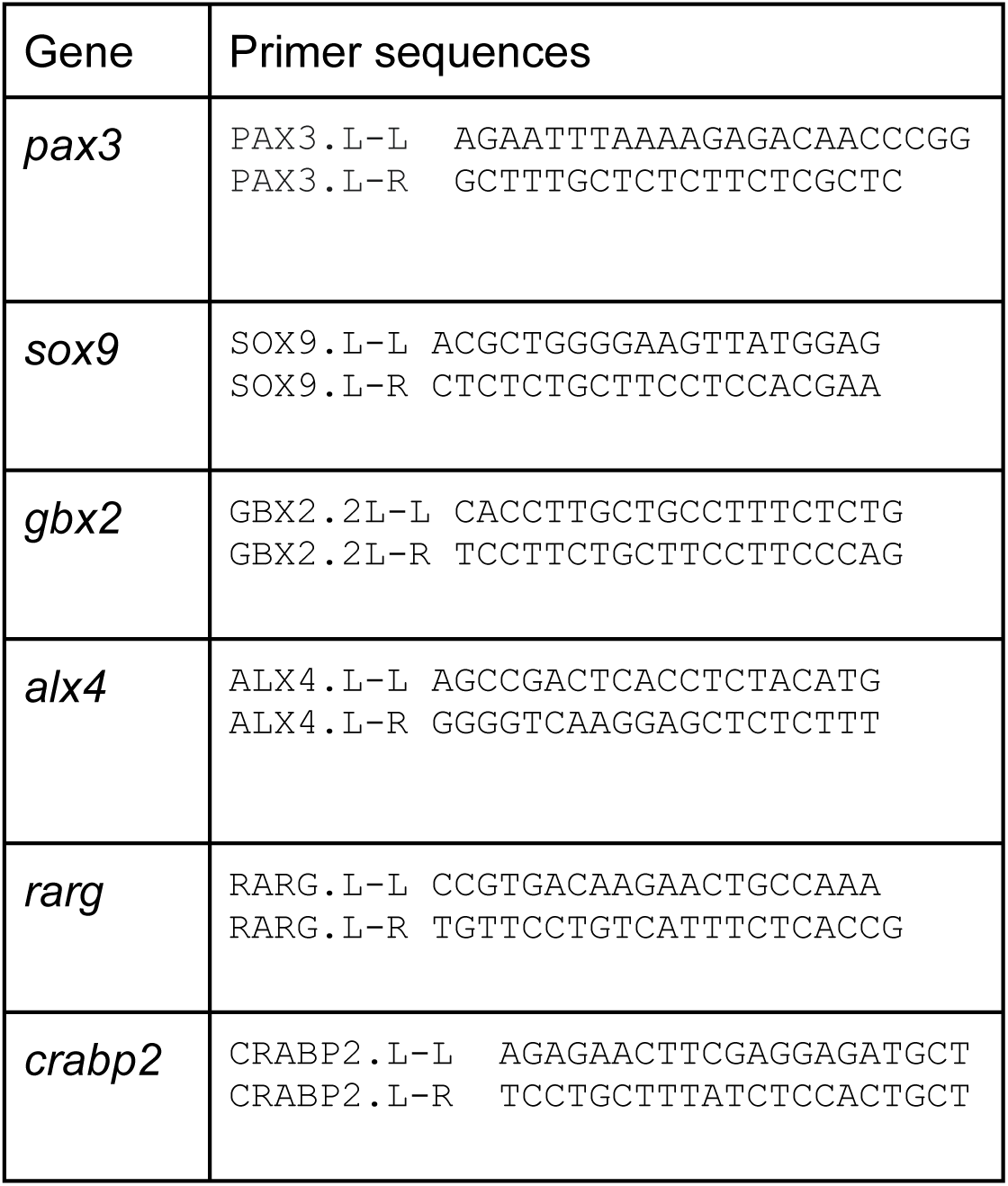
Primer sequences used for RT-qPCR.

